# Biologically Informed Multi-Omics Integration Reveals Clinically Meaningful Patient Representations in Acute Myeloid Leukemia

**DOI:** 10.64898/2026.09.19.752929

**Authors:** Dona Hasini Gammune, Doan Bui, Tongjun Gu

**Affiliations:** Versiti Blood Research Institute, Milwaukee, Wisconsin; Data Science Institute, Medical College of Wisconsin, Milwaukee, Wisconsin; Department of Biostatistics, University of Florida, Gainesville, Florida

**Keywords:** Multi-omics integration, deep learning, graph neural networks, patient representation learning, cross-omics attention, acute myeloid leukemia

## Abstract

Integrating heterogeneous molecular and clinical data into unified, clinically meaningful patient representations remains a major challenge in precision medicine. Here, we present SurvMOCA-GNN, a biologically informed multi-omics integration framework that explicitly incorporates biological network structure, cross-omics regulatory relationships, and patient-specific clinical information to generate unified patient representations. Using gene expression, microRNA expression, and clinical data from the TCGA-LAML cohort, SurvMOCA-GNN revealed patient representations that identified prognostically distinct patient groups and further stratified patients within established European LeukemiaNet 2022 risk categories. Systematic ablation analyses demonstrated that incorporation of graph-based modeling, cross-omics attention, adaptive clinical integration, and biological prior knowledge progressively improved integration quality, resulting in more structured patient representations and stronger prognostic stratification. Compared with existing multi-omics integration approaches, SurvMOCA-GNN consistently produced more biologically coherent patient representations with improved survival discrimination. Evaluation in an independent AML cohort, together with transfer learning analyses, further demonstrated the robustness and transferability of the integrated representations across heterogeneous patient populations. Together, these findings demonstrate that biologically informed multi-omics integration reveals clinically meaningful patient representations and provides a general framework for integrating heterogeneous molecular and clinical data to improve patient stratification in AML and potentially other diseases. The SurvMOCA-GNN framework is freely available at https://github.com/tjgu/SurvMOCA-GNN.git.

## Introduction

Integrating heterogeneous molecular and clinical data remains a fundamental challenge in precision medicine [1,2]. Advances in high-throughput sequencing technologies have enabled comprehensive profiling of multiple biological layers, including gene expression, microRNA (miRNA) expression, epigenetic modifications, and other molecular features [2,3]. These complementary modalities provide distinct yet interconnected perspectives on disease biology and have facilitated important advances in molecular subtyping, biomarker discovery, and disease characterization [3,4]. However, effectively integrating these heterogeneous data remains difficult because of differences in dimensionality, scale, noise characteristics, and modality-specific information content [1,2,5]. Consequently, successful multi-omics integration should generate unified patient representations that preserve complementary biological information, capture disease heterogeneity, and remain robust across independent cohorts [5,6].

Recent machine learning and deep learning approaches have advanced multi-omics integration by enabling nonlinear modeling of high-dimensional biological data and learning low-dimensional patient representations [7–10]. Autoencoders, latent factor models, graph neural networks, and attention-based architectures have shown considerable promise for integrating heterogeneous molecular modalities [6–10]. Despite these advances, many existing methods either treat molecular modalities largely independently or employ generic fusion strategies that do not explicitly incorporate biological network structure, regulatory relationships across molecular layers, or the patient-specific contributions of different data modalities. As a result, the learned patient representations may not fully capture the complementary biological information distributed across molecular modalities, potentially limiting their biological interpretability, robustness, and generalizability across independent cohorts [5,8,10].

Biological systems are inherently organized through coordinated interactions across multiple molecular layers. Among these relationships, miRNAs play a critical role in post-transcriptional regulation of gene expression and contribute to cellular differentiation, proliferation, and therapeutic response [11,12]. Dysregulated miRNA–gene regulatory networks have been implicated in the initiation, progression, and therapeutic resistance of acute myeloid leukemia (AML) [13–15], suggesting that explicitly modeling these interactions may improve the biological relevance of multi-omics integration. Similarly, genes and miRNAs function within interconnected biological networks rather than as independent molecular features, motivating the incorporation of biological structure into multi-omics integration frameworks [16,17].

AML provides an ideal setting for evaluating biologically informed multi-omics integration because of its pronounced molecular and clinical heterogeneity. Despite substantial advances in genomic characterization and the implementation of risk stratification frameworks such as the European LeukemiaNet (ELN)-2022 recommendations [18], considerable variability in clinical outcomes persists within established risk groups, particularly among patients classified as intermediate risk [19,20]. Consequently, patients with similar clinical and genomic profiles can experience markedly different disease trajectories and treatment responses, highlighting limitations of current prognostic models. These observations suggest that clinically relevant disease variation arises from complex interactions across multiple molecular layers and may not be fully captured by existing risk classification frameworks [1,21]. Integrative approaches that explicitly model these multilayer biological relationships may therefore better capture complementary biological information across molecular modalities, leading to more clinically meaningful patient representations and improved patient stratification.

In this study, we developed SurvMOCA-GNN, a biologically informed multi-omics integration framework that integrates gene expression, miRNA expression, and clinical covariates into unified patient representations. By incorporating biological network structure, regulatory interactions across molecular modalities, and patient-specific clinical information, SurvMOCA-GNN is designed to integrate complementary biological information into patient representations that better capture clinically relevant disease heterogeneity. Using AML as a model system, we systematically evaluated the contribution of each architectural component to biologically informed multi-omics integration and assessed whether the resulting patient representations were biologically and clinically meaningful through patient stratification, prognostic analyses, comparisons with existing integration approaches, and external validation.

## Materials and methods

### Problem formulation

Let 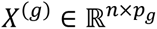 denote gene expression measurements, 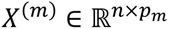 denote miRNA expression measurements, and 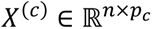 denote clinical covariates for *n* patients. patients. These data modalities provide complementary information describing molecular and clinical variation across patients.

The objective of this study is to learn a latent patient representation (*Z*) that integrates molecular and clinical information while preserving biologically relevant relationships within and across modalities. The learned representation is subsequently used for patient stratification, visualization, and survival analysis.

Formally, we seek to learn a mapping

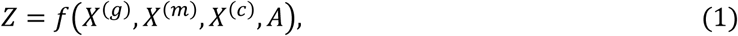

where *A* denotes biological interaction networks constructed from molecular features. Intermediate latent representations generated throughout the framework correspond to modality-specific, graph-refined, and cross-modality-refined embeddings.

### Overview of the Proposed Framework

SurvMOCA-GNN is a multi-omics representation learning framework that integrates gene expression, miRNA expression, and clinical covariates into a unified latent representation (Fig. 1). The framework consists of four sequential stages: modality-specific representation learning, cross-modal interaction modeling, multimodal fusion, and latent representation compression.

**Figure 1.**
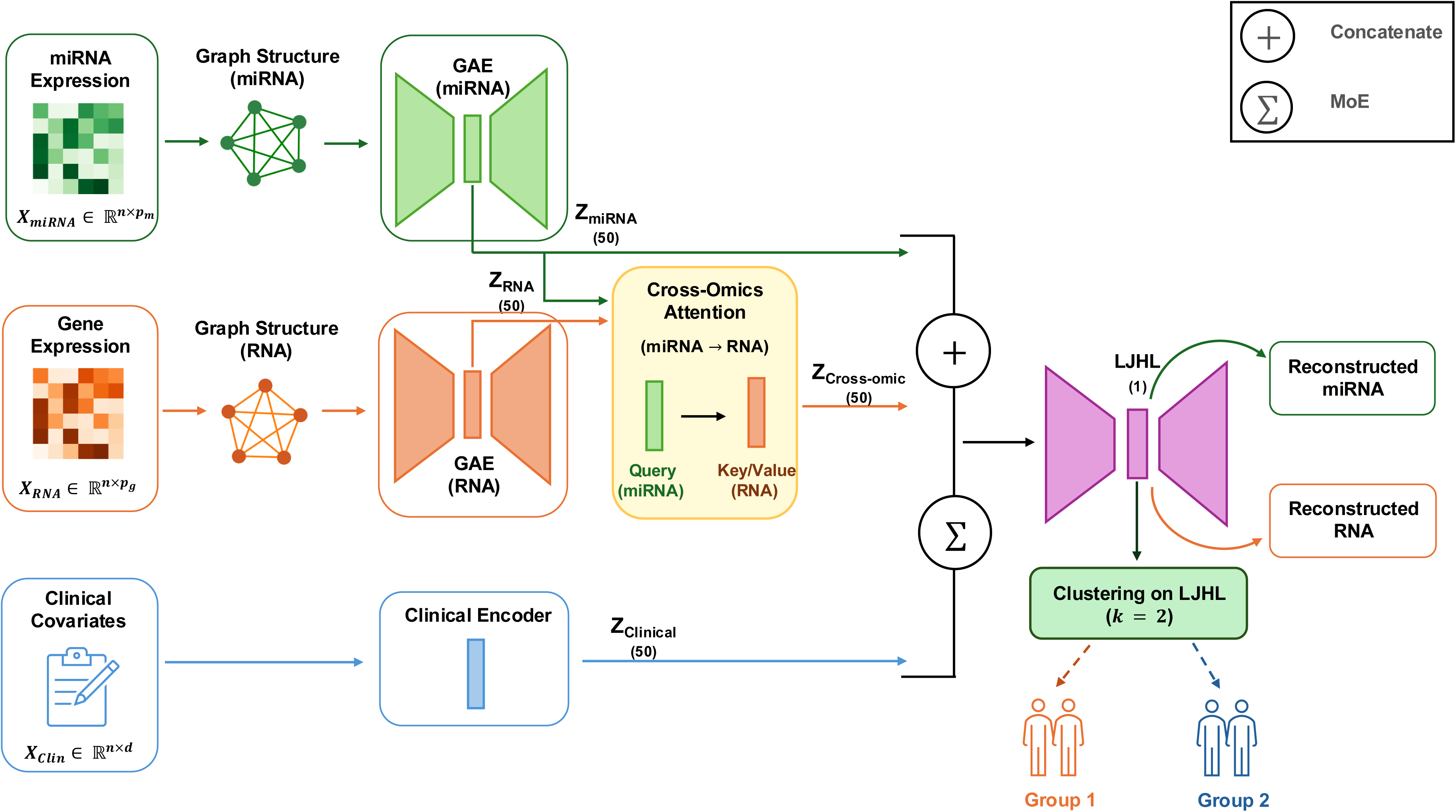
Overview of the SurvMOCA-GNN framework. Schematic illustration of the proposed SurvMOCA-GNN framework for multi-omics integration and survival-associated patient stratification in AML.

Gene expression and miRNA data were first encoded independently using graph autoencoders (GAEs) to generate modality-specific representations while incorporating feature-level network structure. Cross-omics attention was subsequently applied to model interactions between modalities, allowing information transfer between miRNA and gene representations. The resulting embeddings were integrated using a mixture-of-experts (MoE) fusion strategy and combined with clinical covariates. Finally, the fused representation was compressed through a joint autoencoder to obtain a low-dimensional latent representation used for downstream clustering and survival analyses.

The framework was developed to learn patient representations that integrate complementary molecular and clinical information while preserving biological structure across modalities.

### Data and preprocessing

Multi-omics and clinical data were obtained from the Acute Myeloid Leukemia cohort of The Cancer Genome Atlas (TCGA-LAML). Gene expression data were normalized using upper-quartile normalization [22] followed by *log*₂ − transformation. miRNA expression data were normalized using the trimmed mean of M-values (TMM) method [23] and subsequently *log*_2_ − transformed. Low-expression features were removed by retaining genes and miRNAs with counts per million (CPM) greater than 1 in a sufficient number of samples [24]. Additional details regarding preprocessing and quality-control procedures are described in our previous study [25].

To standardize input features across modalities, gene expression and miRNA expression matrices were independently scaled to the range [0,1] using min–max normalization:

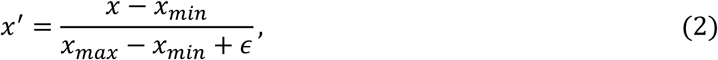

where *x* denotes the original feature value, *x*_min_ and *x*_max_ represent the minimum and maximum values across samples, and *ε* is a small constant introduced to prevent division by zero.

Clinical covariates were preprocessed by one-hot encoding categorical variables and standardizing continuous variables. Samples were aligned across all modalities prior to model training to ensure consistent patient representation.

External validation was performed using an independent AML cohort from the Genomics of Acute Myeloid Leukemia (GAML) study. The trained model was applied to the external cohort to evaluate the robustness and transferability of the learned latent representations.

### SurvMOCA-GNN architecture

The proposed framework consists of four sequential embedding refinement stages: graph-based representation learning, cross-omics attention, adaptive fusion, and clinical covariate integration (Fig. 1). Each stage is designed to enhance the biological and prognostic information contained within the learned patient representation.

### Graph-based embedding refinement

The proposed framework integrates gene expression, miRNA expression, and clinical covariates within a graph-based multi-omics architecture to learn a unified latent representation for AML risk stratification. For each molecular modality, latent features were first learned using GAEs, which capture both expression patterns and underlying biological network structure. Encoder architectures were selected according to the dimensionality and complexity of each molecular modality. Gene expression data were modeled using a deeper architecture than miRNA data because of their substantially larger feature space, while miRNA representations were learned using a more compact encoder [26]. Detailed architectural specifications and hyperparameter settings are provided in the Supplementary Methods.

To improve modality-specific embeddings, gene expression and miRNA data were modeled independently using GAEs [27]. For each modality, a feature-level graph was constructed from molecular correlations, where nodes represent genes or miRNAs and edges represent biological associations between features.

Encoded feature representations were subsequently refined through graph convolutional propagation,

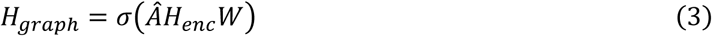

where *H_enc_* denotes encoded feature embeddings, *Â* is the normalized adjacency matrix, and *W* represents learnable parameters [28].

The resulting graph-refined embeddings integrate molecular expression patterns with biological network structure, enabling features with similar biological behavior to occupy nearby regions of the latent space. These graph-enhanced representations serve as the foundation for subsequent cross-omics attention and adaptive fusion stages.

### Cross-omics embedding refinement

While graph-based refinement captures biological structure within each modality, it does not explicitly model regulatory interactions between molecular layers. To address this limitation, we introduced a cross-omics attention mechanism [29] that refines gene embeddings using information derived from miRNA representations. This stage is motivated by the established role of miRNAs in post-transcriptional regulation of gene expression and enables the model to capture context-dependent inter-omic relationships [11,12,30].

Given graph-refined gene and miRNA embeddings, attention is computed using the scaled dot-product formulation:

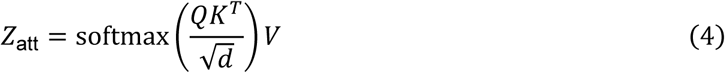

where *Q*, *K*, and *V* denote the query, key, and value representations, respectively, and *d* is the embedding dimension.

The resulting attention-refined representation incorporates regulatory information across modalities and generates more informative patient embeddings than independent modality-specific representations. These refined embeddings are subsequently integrated through the adaptive fusion module.

### Adaptive multi-omics fusion

The graph- and attention-refined modality-specific embeddings were integrated using a MoE fusion strategy [31,32]. Unlike conventional concatenation approaches that assign equal importance to all modalities, the MoE framework adaptively weights modality contributions according to their relevance for representation learning.

The integrated molecular embedding was computed as

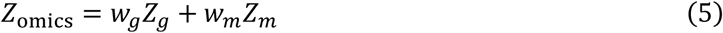

where *Z_g_* and *Z_m_* denote the refined gene and miRNA embeddings, respectively, and *w_g_* and *w_m_* are learned modality-specific weights satisfying *w_g_* + *w_m_* = 1.

This adaptive fusion mechanism enables the model to emphasize the most informative molecular signals while preserving complementary information across modalities. Consequently, the resulting representation provides a more robust and biologically meaningful patient embedding for downstream analysis.

### Clinical embedding augmentation

To incorporate established prognostic factors, clinical covariates, including age, sex, and ELN-2022 risk classification, were encoded and integrated with the molecular representation. Clinical information was incorporated either through direct concatenation or as an additional expert within the MoE framework, enabling adaptive balancing of molecular and clinical signals. The resulting unified representation was used for latent-space learning and downstream survival analyses.

### Final latent representation

The final SurvMOCA-GNN model progressively refines patient representations through graph-based learning, cross-omics attention, adaptive fusion, and clinical augmentation. The resulting integrated representation was compressed through a joint autoencoder with hidden layers of 50 and 10 neurons and a one-dimensional bottleneck layer to generate the final latent embedding.

This embedding represents the culmination of the refinement process and serves as a biologically informed summary of molecular and clinical disease characteristics. The learned latent representation was subsequently used for patient stratification and survival analysis.

### Model training and evaluation

The proposed framework was trained in an unsupervised manner to learn compact and biologically meaningful patient embeddings from multi-omics and clinical data. Training was guided by a joint objective that balances preservation of molecular information with maintenance of biological network structure:

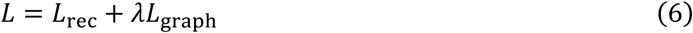

where *L*_rec_ denotes molecular reconstruction loss, *L*_graph_ denotes graph reconstruction loss, and *λ* controls the contribution of graph structure preservation.

Model parameters were optimized using the Adam optimizer [33]. Hyperparameters were selected using Optuna-based optimization [34] with five-fold cross-validation. Candidate configurations were evaluated based on embedding quality, assessed through reconstruction performance and downstream survival stratification. The final model was trained using a learning rate of 0.05, corruption level of 0.2, batch size of 50, and 1000 training epochs. Additional optimization details are provided in the Supplementary Methods.

The primary objective of SurvMOCA-GNN was to generate embeddings that capture clinically relevant disease heterogeneity. Accordingly, model performance was primarily assessed based on survival stratification of patient groups derived from the final latent representation. Secondary analyses evaluated latent-space organization, reconstruction quality, and external generalizability. Comparative performance was further assessed through systematic ablation studies and benchmarking against representative multi-omics integration approaches. Detailed evaluation procedures are provided in the Supplementary Methods.

## Results

### Overview of the SurvMOCA-GNN framework

We developed SurvMOCA-GNN, a biologically informed multi-omics integration framework that integrates gene expression, miRNA expression, and clinical covariates into unified patient representations for AML risk stratification (Fig. 1). The framework combines GAEs [27], cross-omics attention [29], and MoE fusion [31] to capture biological network structure, regulatory interactions between molecular modalities, and patient-specific clinical information. Gene and miRNA features are first encoded using modality-specific GAEs, integrated through cross-omics attention, fused with clinical covariates, and projected into unified patient representations, which are subsequently used to identify patient subgroups by K-means clustering.

### Study cohorts

The proposed framework was trained and evaluated using the TCGA-LAML cohort, comprising 162 patients with matched gene expression, miRNA expression, clinical covariates, and overall survival information. Generalizability was evaluated in the independent GAML cohort, comprising 19 samples from 11 patients (8 AML and 11 matched normal samples). External validation included both direct application of the TCGA-trained model and transfer learning to the GAML cohort. Cohort characteristics are provided in Supplementary Table 1.

### Biologically informed integration preserves complementary molecular information

To assess whether the integrated patient representations preserved information from the original molecular measurements, we evaluated reconstruction performance for both gene expression and miRNA modalities using Pearson correlation and reconstruction loss metrics (Fig. 2a-2b; Supplementary Fig. 1). Reconstruction performance remained comparable between the training and test datasets, indicating that the learned representations preserved molecular information while minimizing overfitting. Comparison of clinical integration strategies showed that direct concatenation achieved slightly higher reconstruction performance than MoE fusion, whereas the MoE framework enabled adaptive weighting of molecular and clinical modalities. These findings indicate that the learned patient representations preserve molecular information while maintaining stable reconstruction performance across the training and test datasets.

**Figure 2.**
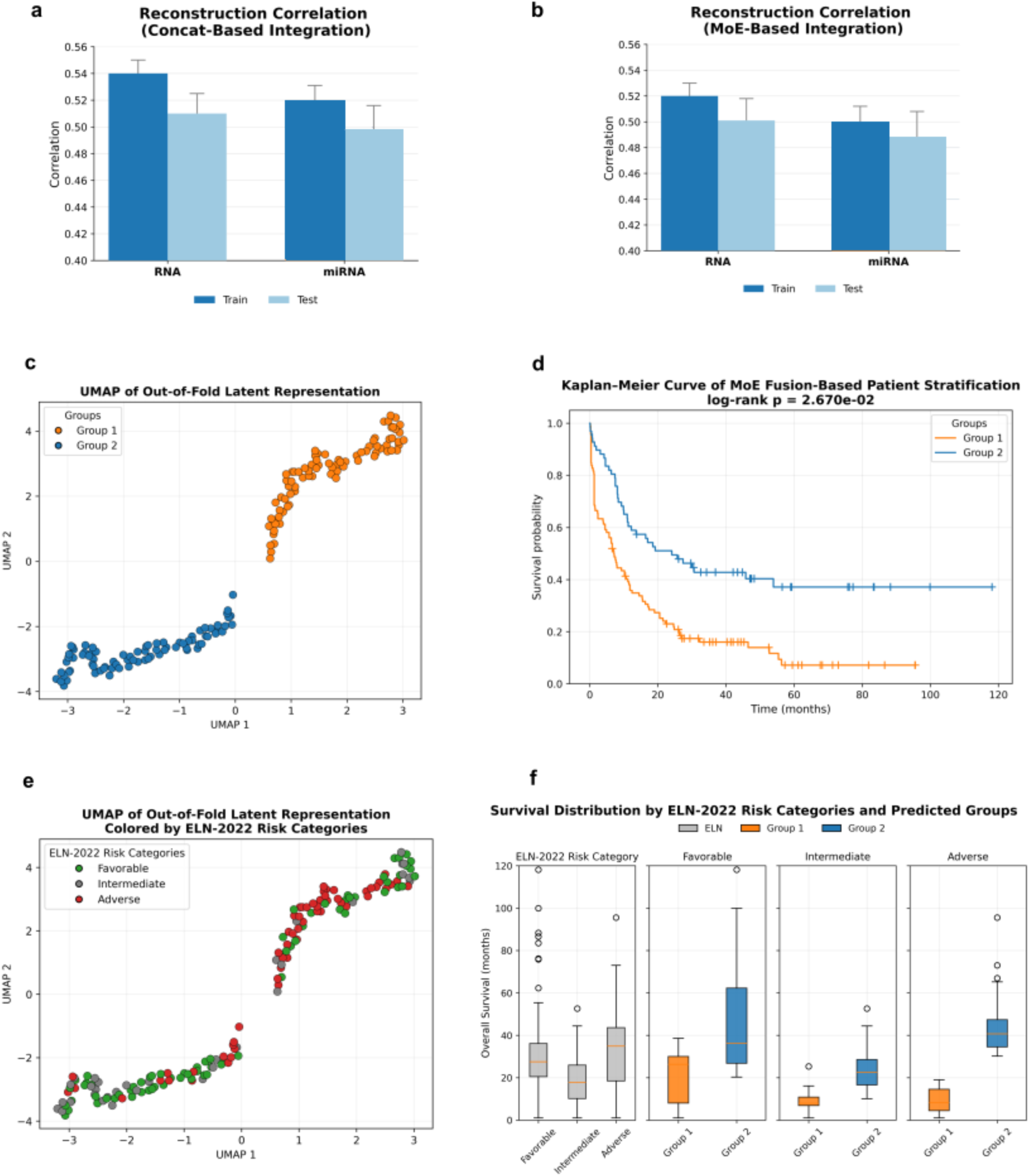
Evaluation of the proposed framework and characterization of latent patient groups. (a) UMAP visualization of out-of-fold latent representations colored by predicted patient groups. (b) Kaplan–Meier survival curves of the predicted patient groups. (c) UMAP visualization of out-of-fold latent representations colored by ELN-2022 risk categories. (d) Distribution of overall survival across ELN-2022 risk categories stratified by predicted patient groups. (e) Reconstruction correlation of the concatenation-based clinical integration model for RNA and miRNA data. (f) Reconstruction correlation of the MoE-based clinical integration model for RNA and miRNA data.

### Integrated patient representations identify prognostically distinct patient groups

To evaluate whether the integrated patient representations captured clinically relevant variation among AML patients, we visualized the out-of-fold representation space using Uniform Manifold Approximation and Projection (UMAP) [35]. The one-dimensional patient representations were partitioned into two groups using K-means clustering (k = 2), and the resulting clusters were projected onto the UMAP for visualization. The learned representation space separated patients into two distinct clusters (Fig. 2c), indicating that the framework learned structured patient representations beyond the original multi-omics feature space.

We next examined whether these representation-derived groups were associated with clinical outcome. Kaplan–Meier analysis [36] demonstrated significantly different overall survival between the two groups (Fig. 2d; *log* − *rank p* = 2.67 × 10^−2^; *Hazard Ratio* (*HR*) = 2.4; 95% *confidence interval*, 1.5 − 4.0). Patients assigned to Group 2 exhibited consistently improved survival compared with those assigned to Group 1, indicating that the learned patient representations capture variation associated with patient outcome.

To determine whether these patterns arose from the learned patient representations rather than the original input features alone, we visualized the raw integrated gene expression and miRNA feature space. In contrast to the learned representation space, the raw feature space exhibited weaker organization and greater overlap between patient groups, accompanied by reduced survival separation (Supplementary Fig. 2). Similar survival trends were observed using the concatenation-based framework (Supplementary Fig. 3). Together, these findings indicate that the integrated patient representations capture clinically relevant variation beyond the original multi-omics feature space, enabling the identification of prognostically distinct patient groups.

### Integrated patient representations provide additional stratification beyond ELN-2022 classification

To assess the relationship between the integrated patient representations and established clinical risk stratification, we visualized the learned representation space according to ELN-2022 risk categories (Fig. 2e). Substantial overlap was observed among ELN-2022 risk categories, particularly within the intermediate-risk group, highlighting the clinical heterogeneity that remains within the current ELN-2022 classification system.

We next examined the distribution of model-derived groups within each ELN-2022 category (Fig. 2f). Within the favorable-risk group, the learned representations identified subpopulations with differing survival patterns, suggesting residual heterogeneity not fully reflected by ELN-2022 classification alone. More pronounced separation was observed within the intermediate-risk group, where patients were divided into subgroups exhibiting distinct survival outcomes. Similar trends were observed within the adverse-risk category, where the learned representations identified subsets of patients with differing prognostic trajectories.

The distinct survival separation observed within each ELN-2022 risk category, with one subgroup exhibiting poorer survival and the other demonstrating improved survival, suggests that the integrated patient representations capture clinically relevant variation not fully reflected by the existing ELN-2022 classification and provide additional resolution of patient heterogeneity.

### Multi-omics integration yields more informative representations than single-modality modeling

To evaluate the contribution of multi-omics integration, we compared SurvMOCA-GNN with models trained using gene expression or miRNA expression alone. Model performance was assessed using reconstruction quality and survival stratification derived from the learned patient representations.

Single-modality models achieved better reconstruction performance for their respective molecular modality but poorer reconstruction performance for the complementary modality (Fig. 3a; Supplementary Fig. 4), indicating that each omics layer contains complementary biological information that cannot be fully captured independently. Accordingly, the integrated framework achieved superior reconstruction performance (Fig. 2a; Supplementary Fig. 1) and stronger survival stratification (Fig. 2d) than either single-modality model (Fig. 3a-3b).

**Figure 3.**
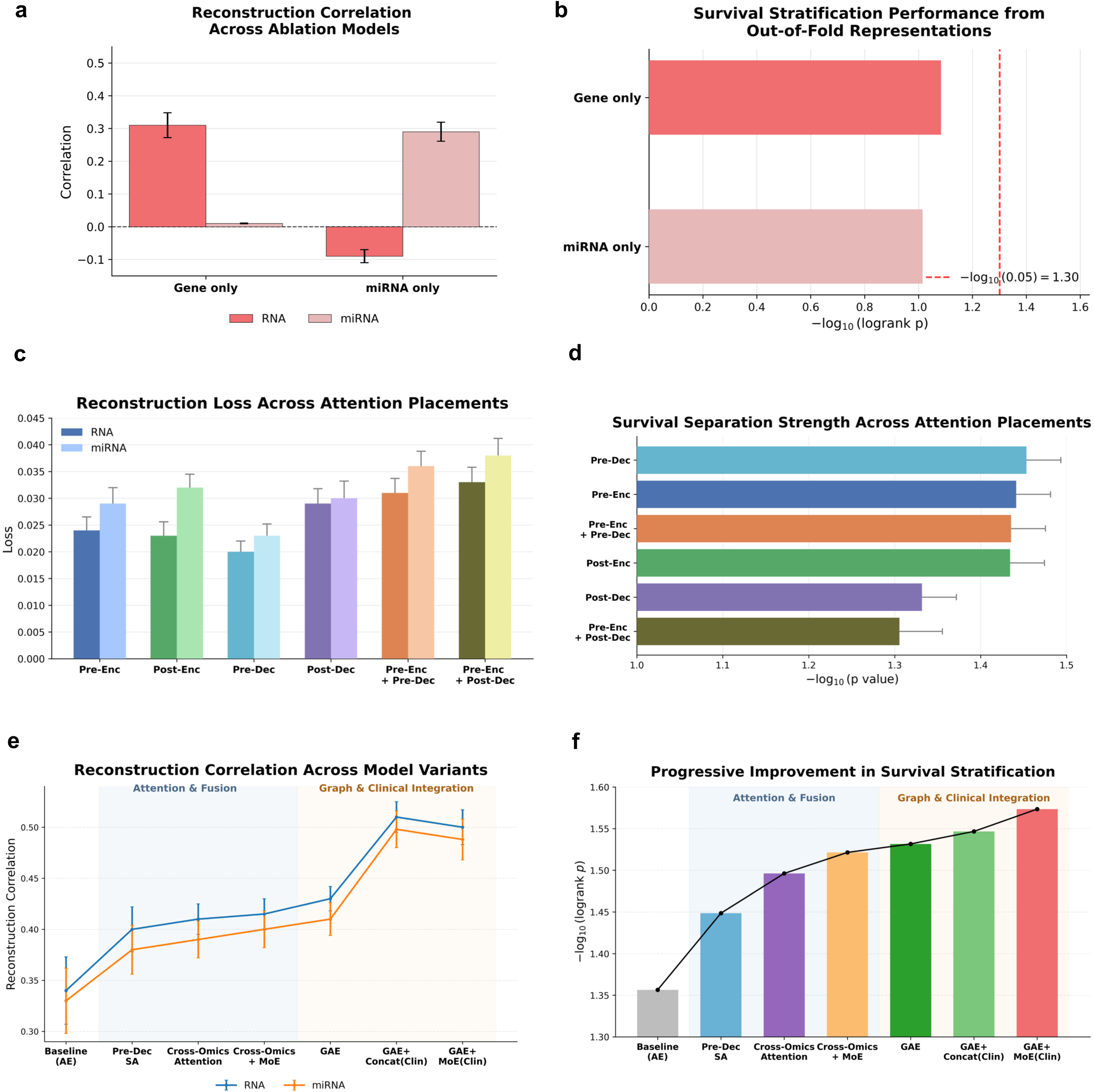
Ablation analysis of the SurvMOCA-GNN architecture. (a) Reconstruction correlation of single-omics models. (b) Survival stratification performance of single-omics models. (c) Reconstruction correlation for different self-attention placements. (d) Survival stratification performance for different self-attention placements. (e) Reconstruction correlation across sequential model variants. (f) Survival stratification performance across sequential model variants.

### Biologically informed integration progressively improves patient representations and prognostic stratification

To evaluate the contribution of individual components of the integration framework, we systematically compared successive model configurations, including direct multi-omics integration, conventional autoencoders (AE), variational autoencoders (VAE) [37], self-attention architectures [32], cross-omics attention, graph autoencoders (GAEs), and alternative clinical covariate integration strategies (Fig. 3c–3f; Supplementary Figs. 5–10).

Direct integration of raw gene expression and miRNA features resulted in weak organization of the learned representation space and limited survival separation (Supplementary Fig. 2). Both AE- and VAE-based models improved representation quality and prognostic separation (Supplementary Fig. 6). However, the AE framework demonstrated more consistent downstream performance and was therefore selected as the foundation for subsequent model development.

We next evaluated multiple self-attention configurations by introducing self-attention at different stages of the model to determine the most effective strategy for modeling intra-modal relationships. Among the evaluated architectures, pre-decoder self-attention consistently achieved the strongest performance across reconstruction and survival analyses (Fig. 3c-3d; Supplementary Figs. 7-8).

Incorporation of cross-omics attention further improved the learned patient representations, and integration of GAEs provided additional improvements. Finally, incorporation of clinical covariates further improved prognostic stratification, with the MoE fusion strategy producing stronger survival separation than simple concatenation, although concatenation achieved slightly better reconstruction performance (Fig. 3e-3f; Supplementary Figs. 9-10).

Collectively, these analyses demonstrate that each biologically motivated component contributes to improved patient representations, resulting in better organization of the representation space, preservation of molecular information, and stronger prognostic stratification.

### SurvMOCA-GNN generates more prognostically informative patient representations than existing multi-omics integration approaches

We compared SurvMOCA-GNN with representative multi-omics integration methods, including MOFA [6], CrossAttOmics [29], and MO-GCAN [38] (Fig. 4a–4d; Supplementary Figs. 11,12). In addition to the original MOFA implementation without clinical information (MOFA-NC), we evaluated a modified version incorporating clinical covariates as an additional modality (MOFA-C) to assess the contribution of clinical information. For all methods, the learned patient representations were subjected to identical downstream clustering and survival analyses to ensure a fair comparison.

**Figure 4.**
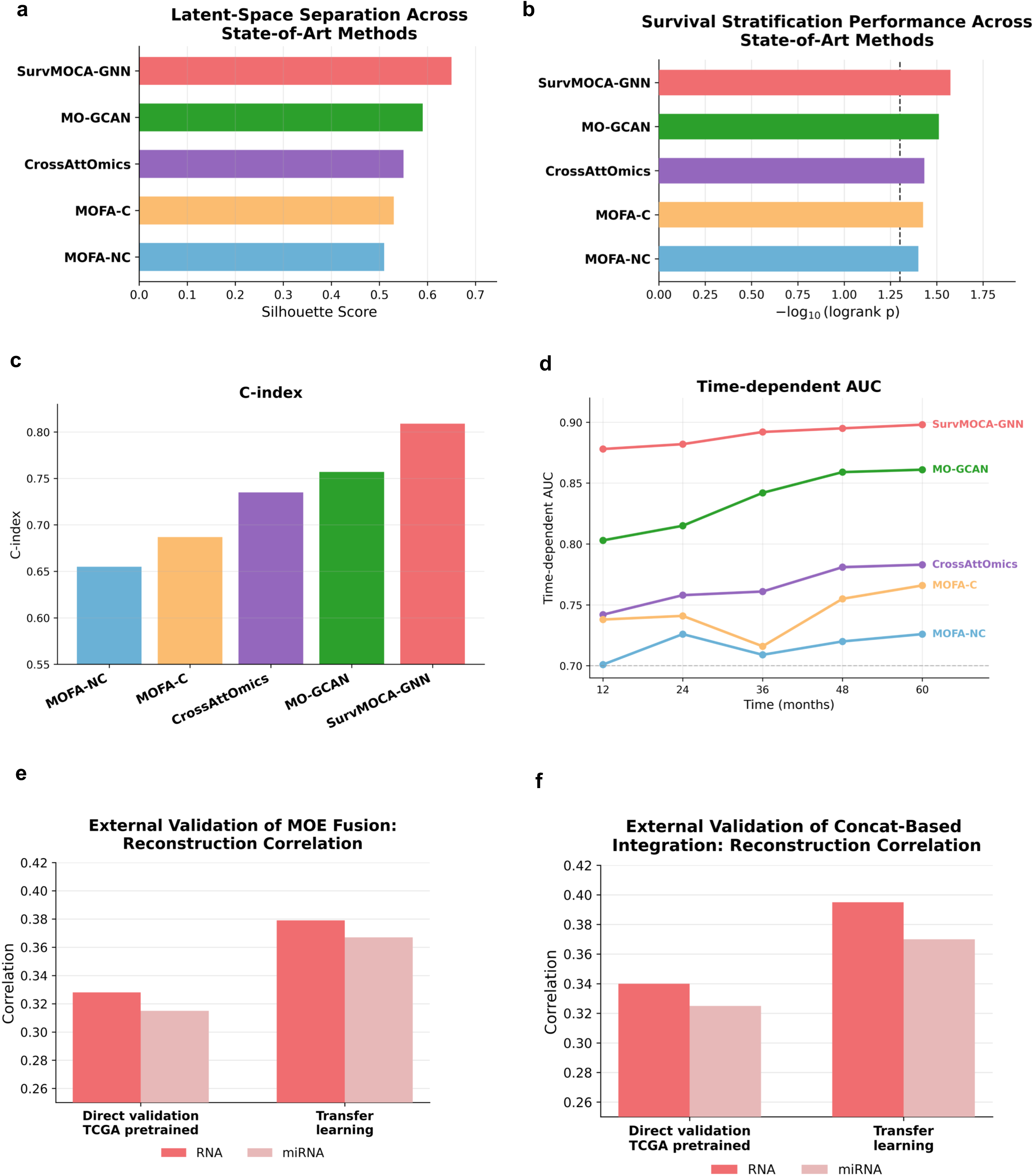
Benchmarking and external validation of SurvMOCA-GNN. (a) Latent-space separation across SurvMOCA-GNN and representative multi-omics integration methods, quantified using silhouette score. (b) Survival stratification performance across methods, evaluated using −log_10_(log-rank p value). (c) Prognostic performance across methods, measured using the C-index. (d) Time-dependent area under the receiver operating characteristic curve (AUC) across multiple follow-up intervals. (e) External validation of the MoE-based integration model on the independent GAML cohort using direct validation and transfer learning. (f) External validation of the concatenation-based integration model on the independent GAML cohort using direct validation and transfer learning.

We first compared the organization of the learned representation space using UMAP visualization and silhouette analysis [39]. SurvMOCA-GNN generated compact and well-separated representation structures (Fig. 2c), achieving the highest silhouette score, which quantifies within-cluster cohesion and between-cluster separation, among the evaluated approaches (Fig. 4a). Progressive improvements were observed from MOFA-NC to MOFA-C, CrossAttOmics, and MO-GCAN, suggesting that incorporation of clinical information, cross-modal relationships, and graph-based modeling progressively improved the organization of the learned representation space (Fig. 4a; Supplementary Fig. 11).

To evaluate the clinical relevance of the learned patient representations, patients were clustered using K-means (k = 2), followed by Kaplan–Meier survival analysis. SurvMOCA-GNN produced the strongest survival separation among the evaluated methods (Fig. 4b; Supplementary Fig. 12). Incorporation of clinical covariates improved the performance of MOFA, and both CrossAttOmics and MO-GCAN demonstrated stronger prognostic separation than the latent factor-based approaches. Nevertheless, SurvMOCA-GNN consistently generated patient groups with the greatest survival separation. Similar trends were observed in the hazard ratio analysis, where SurvMOCA-GNN achieved the largest separation between the identified risk groups (Supplementary Fig. 12).

We next evaluated whether improved patient representations were associated with enhanced prognostic performance. SurvMOCA-GNN achieved the highest concordance index (C-index) [40] among all evaluated methods (Fig. 4c). Consistent with this finding, time-dependent area under the receiver operating characteristic curve (AUC) analysis demonstrated superior predictive performance across all evaluated follow-up intervals (Fig. 4d).

Collectively, these findings demonstrate that SurvMOCA-GNN consistently outperformed existing multi-omics integration approaches in representation quality, patient stratification, and prognostic prediction.

### Integrated patient representations are transferable across an independent AML cohort

To evaluate the generalizability of SurvMOCA-GNN beyond the discovery cohort, we applied the framework to an independent AML dataset (GAML) and assessed the learned patient representations using reconstruction performance, representation-space visualization, and transfer learning analyses (Fig. 4e–4f; Supplementary Figs. 13-14). Because survival annotations and ELN-2022 risk classifications were unavailable for this cohort, evaluation focused on reconstruction performance, preservation of representation structure, and concordance with biological phenotype.

Both the MoE and concatenation-based integration strategies maintained stable reconstruction performance in the external cohort despite differences in cohort composition and molecular profiles, with performance comparable to that observed in the discovery cohort (Fig. 4e–4f; Supplementary Fig. 13). Direct application of the pretrained model also preserved the overall representation structure, producing two well-separated groups that showed substantial concordance with tumor and normal status (Supplementary Fig. 14). Notably, the few normal samples located within the tumor-associated cluster originated from the same patient as neighboring tumor samples, suggesting that the learned patient representations preserve patient-specific molecular characteristics alongside disease-associated variation.

To further evaluate transferability, we initialized the framework using representations learned from the discovery cohort and subsequently adapted the model to the external dataset through transfer learning. Transfer learning further improved reconstruction performance relative to direct application of the pretrained model (Fig. 4e–4f). The adapted patient representations also exhibited more compact and better-separated clusters, with stronger concordance between the model-derived groups and tumor/normal labels (Supplementary Fig. 14). Similar to the direct application results, the remaining mixed samples corresponded to paired tumor and normal samples from the same patient, further supporting preservation of patient-specific molecular characteristics within the learned representation space.

Together, these findings demonstrate that the patient representations learned by SurvMOCA-GNN are transferable across independent cohorts and can be further refined through transfer learning, supporting the robustness and generalizability of the proposed biologically informed multi-omics integration framework.

## Discussion

In this study, we developed SurvMOCA-GNN, a biologically informed multi-omics integration framework that integrates biological network structure, cross-omics regulatory relationships, and clinical information to generate unified patient representations. Using AML as a model system, we demonstrated that biologically informed integration revealed clinically meaningful patient representations associated with improved prognostic stratification, additional resolution within ELN-2022 risk groups, and robust performance across independent cohorts. Collectively, these findings suggest that explicitly incorporating biological knowledge into multi-omics integration provides a principled strategy for generating clinically informative patient representations.

A central finding of this study is that incorporating biological knowledge throughout the integration process progressively improved the resulting patient representations. Rather than relying solely on statistical relationships within high-dimensional molecular data, SurvMOCA-GNN integrates complementary sources of biological information, including molecular interaction networks, miRNA–gene regulatory relationships, and patient-specific clinical information. Together, these biological relationships guide the integration of heterogeneous molecular modalities, enabling the resulting patient representations to better capture coordinated biological variation associated with disease heterogeneity. These findings suggest that incorporating prior biological knowledge throughout the integration process facilitates biologically informed integration by preserving molecular relationships that may otherwise be overlooked by purely data-driven approaches.

Our findings further suggest that the value of multi-omics integration extends beyond combining heterogeneous molecular datasets. Instead, its primary advantage lies in integrating complementary biological information distributed across different molecular modalities into unified patient representations. The superior performance of the integrated gene expression and miRNA framework compared with single-modality models supports the view that AML heterogeneity arises from coordinated dysregulation across multiple molecular layers rather than any individual omics profile. These findings highlight the importance of integrating complementary molecular information for characterizing disease heterogeneity and improving patient stratification.

Despite major advances in molecular characterization and the widespread adoption of the ELN-2022 recommendations [18], substantial variability in clinical outcomes remains within established risk categories, particularly among patients classified as intermediate risk. The ability of SurvMOCA-GNN to identify prognostically distinct subgroups within existing ELN-2022 categories suggests that clinically relevant biological variation remains incompletely captured by current risk stratification frameworks. Rather than replacing established clinical classification systems, biologically informed multi-omics integration may complement them by revealing additional layers of disease heterogeneity and supporting more refined patient stratification.

Comparison with existing multi-omics integration approaches suggests that the benefits of biologically informed integration extend beyond improvements in predictive performance alone. Although statistical factorization and feature-fusion methods effectively integrate heterogeneous molecular datasets, our findings indicate that explicitly incorporating biological organization— including molecular interaction networks, cross-omics regulatory relationships, and clinical information—provides additional value for characterizing disease heterogeneity. More broadly, these results suggest that the effectiveness of multi-omics integration depends not only on the underlying learning architecture but also on how biological knowledge is incorporated into the integration process.

The ability of SurvMOCA-GNN to generalize across an independent AML cohort further supports the robustness of the proposed integration strategy. Despite differences in cohort composition and molecular characteristics, the framework maintained consistent performance following direct application, while transfer learning further improved adaptation to the external dataset. These findings suggest that biologically informed multi-omics integration can capture transferable molecular patterns that extend beyond the discovery cohort rather than overfitting to dataset-specific characteristics. Such robustness is particularly important for future translational applications, where computational models must remain reliable across diverse patient populations, sequencing platforms, and clinical settings.

Several limitations of this study should be acknowledged. First, although SurvMOCA-GNN was evaluated using an independent AML cohort, further validation in larger, multi-center cohorts and prospective studies will be important to establish its robustness and clinical utility. Second, the current framework integrates gene expression, miRNA expression, and clinical information; incorporating additional molecular modalities, such as DNA methylation, chromatin accessibility, or proteomic data, may further enhance its ability to characterize disease heterogeneity. Finally, although the framework demonstrated consistent prognostic performance, improving model interpretability remains an important direction for future work to better understand the biological mechanisms underlying patient stratification and facilitate clinical adoption.

In conclusion, SurvMOCA-GNN demonstrates that explicitly incorporating biological knowledge into multi-omics integration provides an effective strategy for generating clinically meaningful patient representations. By integrating biological network structure, cross-omics regulatory relationships, and clinical information, the framework improved patient stratification, provided additional resolution within established ELN-2022 risk groups, and generalized across independent cohorts. More broadly, our findings suggest that biologically informed integration offers a general framework for integrating heterogeneous molecular and clinical data that is applicable beyond AML and may support precision oncology in other complex diseases.

## Key Points

- We developed SurvMOCA-GNN, a biologically informed graph neural network framework for integrating multi-omics and clinical data into unified patient representations.
- The framework combines molecular network structure, cross-omics regulatory attention, and adaptive mixture-of-experts fusion to model complementary information across gene expression, miRNA expression, and clinical data.
- Systematic ablation analyses demonstrate that progressively incorporating biological and clinical information improves representation structure and prognostic stratification.
- SurvMOCA-GNN identifies clinically meaningful patient subgroups, including additional prognostic heterogeneity within ELN-2022 risk categories, and outperforms representative multi-omics integration approaches.
- Independent-cohort evaluation and transfer learning demonstrate the robustness and transferability of the learned patient representations.

## Acknowledgments

We thank Drs. Timothy J. Ley and Christopher A. Miller (Washington University); Richard Corbett, and Drs. Emilia Lim and Marco Marra (University of British Columbia) for their invaluable assistance in acquiring the gene and miRNA datasets. This work was supported, in part, by Institutional Research Grant IRG #22-151-37-IRG from the American Cancer Society and by the Medical College of Wisconsin (MCW) Cancer Center.

## Authors’ Contributions

T.G. conceived and supervised the study. D.B. acquired and preprocessed the data. D.H.G. performed the analyses. T.G. and D.H.G. interpreted the results and wrote the manuscript. All authors reviewed, edited, and approved the final version.

## Competing Interests

The authors declare no competing financial interests.

## Data Availability Statement

TCGA-LAML RNA- and miRNA- sequencing data, along with clinical annotations, are publicly available via the NCI Genomic Data Commons (Project ID: TCGA-LAML, Program: TCGA). GAML RNA/small RNA-sequencing data are available via dbGaP (accession ID: phs000159).

## Notes

The code is available at https://github.com/tjgu/SurvMOCA-GNN.git.

## Notes

### Competing Interest Statement

The authors have declared no competing interest.

