## Supplementary Methods, Tables and Figures for "Biologically Informed Multi-Omics Integration Reveals Clinically Meaningful Patient Representations in Acute Myeloid Leukemia"


**Supplementary Methods.**………………………………………………………………………………… 2-9

**Supplementary Tables**

Table 1. Characteristics of the AML cohorts and multi-omics datasets used for model development and external validation……………………………………………………………………………... 10

**Supplementary Figures**

Figure 1. Reconstruction loss of the SurvMoCA-GNN framework with clinical covariate integration… 11

Figure 2. UMAP visualization and survival analysis using raw multi-omics input data………………… 12

Figure 3. Kaplan–Meier survival analysis of the concatenation-based clinical integration model……. 13

Figure 4. Cross-modality reconstruction loss for single-omics models……………………………...... 14

Figure 5. Baseline multi-omics autoencoder model architecture ……………………………………… 15

Figure 6. Comparison of autoencoder and variational autoencoder models.…………………………… 16

Figure 7. Self-attention placement strategies evaluated…………………………..…………………... 17

Figure 8. Reconstruction loss across self-attention placements ………………………………………. 18

Figure 9. Reconstruction loss following the incremental incorporation of architectural components.… 19

Figure 10. Hazard ratios following the incremental incorporation of architectural components……… 20

Figure 11. Comparison of latent-space organization across multi-omics integration methods………. 21

Figure 12. Comparison of survival stratification performance across multi-omics integration methods... 22

Figure 13. External validation reconstruction loss of RNA and miRNA modalities on the external GAML cohort using direct validation of TCGA-trained models and transfer learning………... 23

Figure 14. Generalizability of learned latent representations in the independent GAML cohort……… 24

**Supplementary References**………………………………………………………………………………… 25

### Supplementary Methods

#### Graph Construction and Graph Autoencoder Formulation

For each molecular modality, a feature-level graph was constructed in which nodes represent molecular features (genes or miRNAs) and edges represent co-expression relationships derived from patient expression profiles.

##### *Graph construction*

Let

$$X_g \in \mathbb{R}^{n \times p_g}$$

and

$$X_m \in \mathbb{R}^{n \times p_m}$$

denote the gene expression and miRNA expression matrices, respectively, where $n$ is the number of patients, $p_g$ is the number of genes, and $p_m$ is the number of miRNAs. For each modality, we construct a graph $G = (V, E)$, where nodes represent molecular features (genes or miRNAs) and edges encode relationships derived from their expression profiles across patients.

Edges are constructed based on pairwise similarity between features using Pearson correlation, following established approaches for gene co-expression network construction [1]. For each feature $i$, correlation coefficients with all other features $j$ are computed as:

$$C_{ij} = \mathrm{corr}(X_i, X_j),$$

forming a similarity matrix over the feature space. Self-correlations are excluded by setting $C_{ii} = 0$. To retain only strong associations, edges are defined using a threshold $\tau$:

$$A_{ij} = \begin{cases} | C_{ij} |, & \text{if } | C_{ij} | > \tau, \\ 0, & \text{otherwise.} \end{cases}$$

The adjacency matrix is symmetrized to ensure an undirected graph:

$$A = \max(A, A^{\top}).$$

Finally, the adjacency matrix is row-normalized to obtain the propagation matrix:

$$\tilde{A}_{ij} = \frac{A_{ij}}{\sum_j A_{ij}}.$$

This construction preserves high-confidence relationships while ensuring a stable propagation structure.

***Graph autoencoder***

Given the input data, modality-specific encoder networks first produce low-dimensional feature representations:

$$H_g = f_g(X_g), \qquad H_m = f_m(X_m),$$

where $H_g$ and $H_m$ denote encoded gene and miRNA features, respectively. For gene expression, the encoder consists of two fully connected layers (500 and 50 units) according to our previous work [2], while for miRNA, a single hidden layer with 50 units is used.

In the graph formulation, each node corresponds to a molecular feature (gene or miRNA), and the associated node features are given by the encoded representations $H$. Specifically, each row of $H_g$ represents the latent feature vector for a gene, learned from its expression profile across patients.

These encoded feature representations serve as the input to the graph autoencoder, where graph-based propagation is applied using the normalized adjacency matrix [3]. For gene expression:

$$Z_g = \sigma(\bar{A}_g H_g W_g),$$

and similarly for miRNA:

$$Z_m = \sigma(\bar{A}_m H_m W_m),$$

where $W_g$ and $W_m$ are learnable weight matrices and $\sigma(\cdot)$ denotes a nonlinear activation function.

To preserve graph structure, the adjacency matrices are reconstructed using an inner-product decoder [4]:

$$\hat{A}_g = \sigma\left(Z_g \left(Z_g\right)^T\right), \quad \hat{A}_m = \sigma(Z_m (Z_m)^T).$$

This reconstruction objective encourages features connected in the original graph to remain close in the embedding space.

The learned feature-level embeddings are then projected back to the patient space:

$$\bar{Z}_g = X_g Z_g, \quad \bar{Z}_m = X_m Z_m.$$

This projection integrates observed molecular measurements with graph-derived structural information, producing biologically informed patient representations for downstream multi-omics integration and survival analysis [5,6].

By combining nonlinear feature transformation with graph-based encoding and reconstruction, the proposed GAE framework captures intra-omic biological dependencies and produces robust, biologically meaningful representations for downstream multi-omics integration and survival analysis.

### Cross-Omics Attention

#### Query–key–value projections

To model interactions between molecular modalities, a cross-attention mechanism was applied between graph-refined gene and miRNA representations [7]. Given graph-refined representations $Z^{(m)}$ and $Z^{(g)}$, the attention-refined gene representation is computed as:

$$Z_{\text{att}} = \text{CrossAtt}(Z_m \rightarrow Z_g).$$

This operation follows the standard query–key–value attention formulation, where the target modality (gene) provides the queries and the source modality (miRNA) provides the keys and values:

$$Q = Z_g W_Q, \qquad K = Z_m W_K, \qquad V = Z_m W_V,$$

where $W_Q$, $W_K$, and $W_V$ are learnable projection matrices.

#### Cross-attention operation

The attention-refined representation was computed using scaled dot-product attention:

$$Z_{\text{att}} = \text{softmax}\left(\frac{QK^T}{\sqrt{d_k}}\right)V,$$

where $d_k$ denotes the key dimension.

This formulation allows each gene representation to selectively incorporate information from miRNA representations, thereby capturing learned miRNA–gene associations in the latent representation space. These learned associations are interpreted in the context of the established biological role of miRNAs in regulating gene expression [8,9].

#### Attention placement analysis

To determine the most effective stage for attention-based feature refinement, self-attention (SA) [10] modules were evaluated at multiple locations within the modality-specific autoencoder architecture. Six configurations were considered (Supplementary Fig. 7):

1. Attention before encoder
2. Attention after encoder
3. Attention before decoder
4. Attention after decoder
5. Attention before encoder and before decoder

6. Attention before encoder and after decoder

Model configurations were compared using reconstruction correlation, reconstruction loss, and downstream survival stratification performance. The configuration that yielded the strongest overall performance is adopted in the final model.

### Adaptive Multi-Omics Fusion

#### Molecular Fusion Using a Mixture-of-Experts Framework

Following graph-based encoding and cross-omics attention, gene and miRNA embeddings were integrated using a Mixture-of-Experts (MoE) [11,12] framework. Let $Z_g$ and $Z_m$ denote the gene and miRNA embeddings, respectively. The fused molecular representation was computed as

$$Z_{\text{omics}} = \sum_{i=1}^{M} w_i \, Z_i,$$

where $M = 2$ denotes the number of molecular modalities and $w_i$ represents the learned modality weight.

The modality weights satisfy

$$\sum_{i=1}^{M} w_i = 1, \qquad w_i \geq 0, \forall i.$$

Weights were generated through a softmax-based gating network:

$$s_i = f_{\text{gate}}(Z_i),$$
$$w_i = \frac{\exp(s_i)}{\sum_{j=1}^{M} \exp(s_j)},$$

where $s_i$ denotes the gating score for modality $i$.

The resulting representation $Z_{\text{omics}}$ was used as the molecular embedding for downstream integration.

### Clinical Covariate Integration

#### Concatenation-Based Clinical Integration

As an initial strategy, clinical covariates were encoded into a latent representation $Z_c$ and concatenated with the molecular embedding:

$$Z_{\text{concat}} = [Z_{\text{omics}} \parallel Z_c],$$

where ‖ denotes vector concatenation.

#### MoE-Based Clinical Integration

To enable adaptive weighting of molecular and clinical information, clinical covariates were incorporated as an additional expert within the MoE framework. The final fused representation was computed as

$$Z_{\text{fusion}} = \sum_{i=1}^{M} w_i Z_i,$$

where,

$$Z_i \in \{Z_g, Z_m, Z_c\},$$

and $M = 3$.

The modality weights were determined using the same softmax gating mechanism. The resulting representation $Z_{\text{fusion}}$ was subsequently used for joint latent representation learning and survival-associated patient stratification.

#### Joint Latent Representation Learning

The fused multi-modal representation was further compressed using a joint autoencoder to learn a compact patient-level latent representation. The encoder consisted of two fully connected hidden layers containing 50 and 10 neurons, followed by a one-dimensional bottleneck layer:

$$Z_{\text{fusion}} \to 50 \to 10 \to 1.$$

The one-dimensional bottleneck variable,

$$z \in \mathbb{R},$$

represents the final latent embedding learned by the model. This latent representation was subsequently used for patient clustering, visualization, and survival analyses.

#### Loss Functions

#### Molecular Reconstruction Loss

To preserve modality-specific molecular information, reconstruction loss was computed as the sum of gene and miRNA reconstruction errors:

$$L_{\text{rec}} = \| X_g - \hat{X}_g \|_2^2 + \| X_m - \hat{X}_m \|_2^2,$$

where $X_g$ and $X_m$ denote the original gene and miRNA expression matrices, respectively, and $\hat{X}_g$ and $\hat{X}_m$ represent their reconstructed counterparts.

**Graph Reconstruction Loss**

To preserve biological network structure, graph reconstruction loss was defined as

$$L_{\text{graph}} = \| A_g - \hat{A}_g \|_2^2 + \| A_m - \hat{A}_m \|_2^2,$$

where $A_g$ and $A_m$ denote the original adjacency matrices and $\hat{A}_g$and $\hat{A}_m$denote the reconstructed adjacency matrices for the gene and miRNA graphs, respectively.

**Total Loss**

The overall training objective was formulated as

$$L = L_{\text{rec}} + \lambda L_{\text{graph}},$$

where $\lambda$ controls the contribution of graph structure preservation relative to molecular reconstruction.

**Hyperparameter optimization**

Hyperparameters, including learning rate, corruption level, batch size, and number of training epochs, were optimized using the Optuna framework [13]. The search space included learning rates $\{0.005, 0.01, 0.05, 0.1, 0.15, 0.2, 0.3, 0.5\}$, corruption levels $\{0.01, 0.05, 0.1, 0.2, 0.5\}$, batch sizes $\{10, 20, 50, 100\}$, and training epochs $\{1000, 2000, 5000\}$.

Model performance during optimization was evaluated using five-fold cross-validation. Candidate configurations were assessed using reconstruction quality and downstream embedding performance, and the final model was selected based on consistent performance across validation folds.

The optimal configuration consisted of a learning rate of $0.05$, corruption level of $0.2$, batch size of $50$, and $1000$ training epochs, and was used for all subsequent analyses.

**Evaluation strategy**

To comprehensively evaluate the learned embeddings, we assessed reconstruction quality, latent-space organization, survival stratification, and external generalizability.

**Reconstruction performance**

Reconstruction quality was quantified using Pearson correlation between original and reconstructed molecular profiles for each modality. Reconstruction error was additionally measured using mean squared error (MSE). These metrics assess the extent to which biologically relevant molecular information is preserved within the learned embeddings.

**Clustering, visualization, and survival analysis**

The final latent representation obtained from the one-dimensional bottleneck layer was used for patient stratification. K-means clustering ($k = 2$) was applied to identify patient subgroups representing distinct molecular risk profiles.

To visualize latent-space organization and assess cluster separation, Uniform Manifold Approximation and Projection (UMAP) [14] was applied to the learned embeddings. The clinical relevance of the identified patient groups was evaluated using Kaplan–Meier survival analysis [15], with statistical significance assessed using the log-rank test. Survival stratification served as the primary measure of embedding quality throughout the study.

**Ablation Studies**

A progressive ablation analysis was performed to quantify the contribution of each component of the proposed framework. Starting from a baseline AE, model complexity was incrementally increased through the addition of a VAE [16], self-attention, cross-omics attention, MoE fusion, GAE refinement, and clinical covariate integration. At each stage, model performance was evaluated using reconstruction quality, latent-space organization, and downstream survival stratification.

**Benchmarking and External Validation**

**Benchmarking**

The proposed SurvMOCA-GNN framework was benchmarked against representative multi-omics integration approaches, including MOFA [17], CrossAttOmics [6], and MO-GCAN [18]. The quality and clinical relevance of the learned latent representations were evaluated using complementary metrics. Latent-space organization was assessed using UMAP visualization and quantified using the silhouette score. The ability of the embeddings to capture survival-associated patient heterogeneity was evaluated through Kaplan–Meier analysis, log-rank testing, concordance index (C-index), and time-dependent area under the receiver operating characteristic curve (AUC). Together, these metrics assess the extent to which the learned embeddings preserve biologically and clinically meaningful structure relevant to patient outcomes. To ensure a fair comparison, latent representations generated by each method were subjected to an identical downstream analysis pipeline consisting of dimensionality reduction, clustering, and survival analysis.

**External Validation**

Model generalizability was evaluated using the independent GAML cohort. The trained model was applied to external samples to assess the stability and transferability of the learned latent representations across cohorts.

**Transfer Learning**

To further evaluate cross-cohort robustness, transfer learning experiments were performed by fine-tuning the TCGA-trained model on GAML samples. Performance before and after fine-tuning was compared to assess the extent to which learned molecular representations could be adapted to an independent dataset.

**Supplementary Table 1.**

**Characteristics of the AML cohorts and multi-omics datasets used for model development and external validation.**

| **Characteristic** | **TCGA-LAML** | **GAML** |
|---|---|---|
| Total samples | 162 | 19 |
| AML | 162 | 8 |
| Normal controls | — | 11 |
| Gene expression features | 14265 | 45696 |
| miRNA features | 763 | 707 |
| Clinical covariates | Age, Sex, ELN-2022 risk categories | Sex |
| Survival data | Yes | No |
| Purpose | Model development | External validation |

**Supplementary Fig 1.**

**Reconstruction loss of the SurvMoCA-GNN framework with clinical covariate integration.**

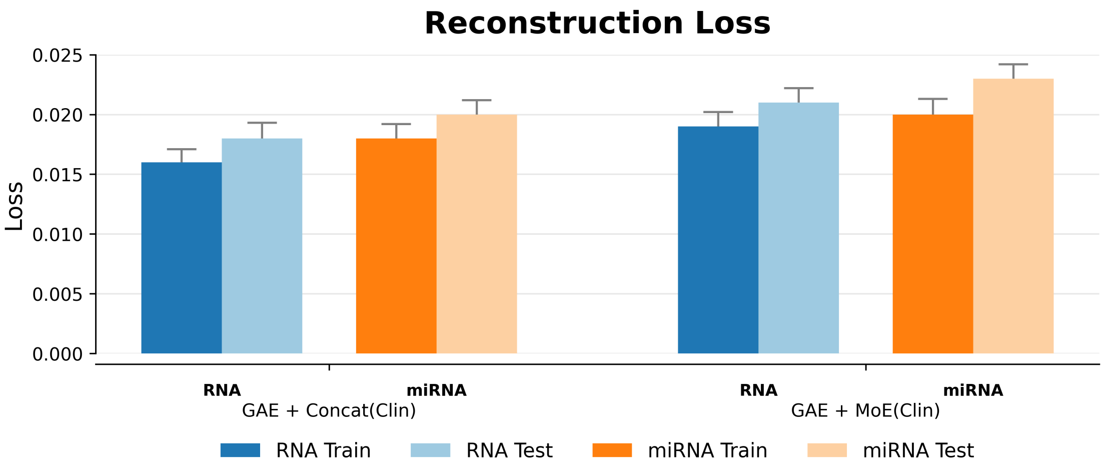

**Supplementary Fig 2.**

**UMAP visualization and survival analysis using raw multi-omics input data.**

(a) UMAP visualization of the raw integrated gene expression and miRNA expression data. (b) Kaplan–Meier survival analysis of the two patient groups identified by K-means clustering from the raw integrated molecular features.

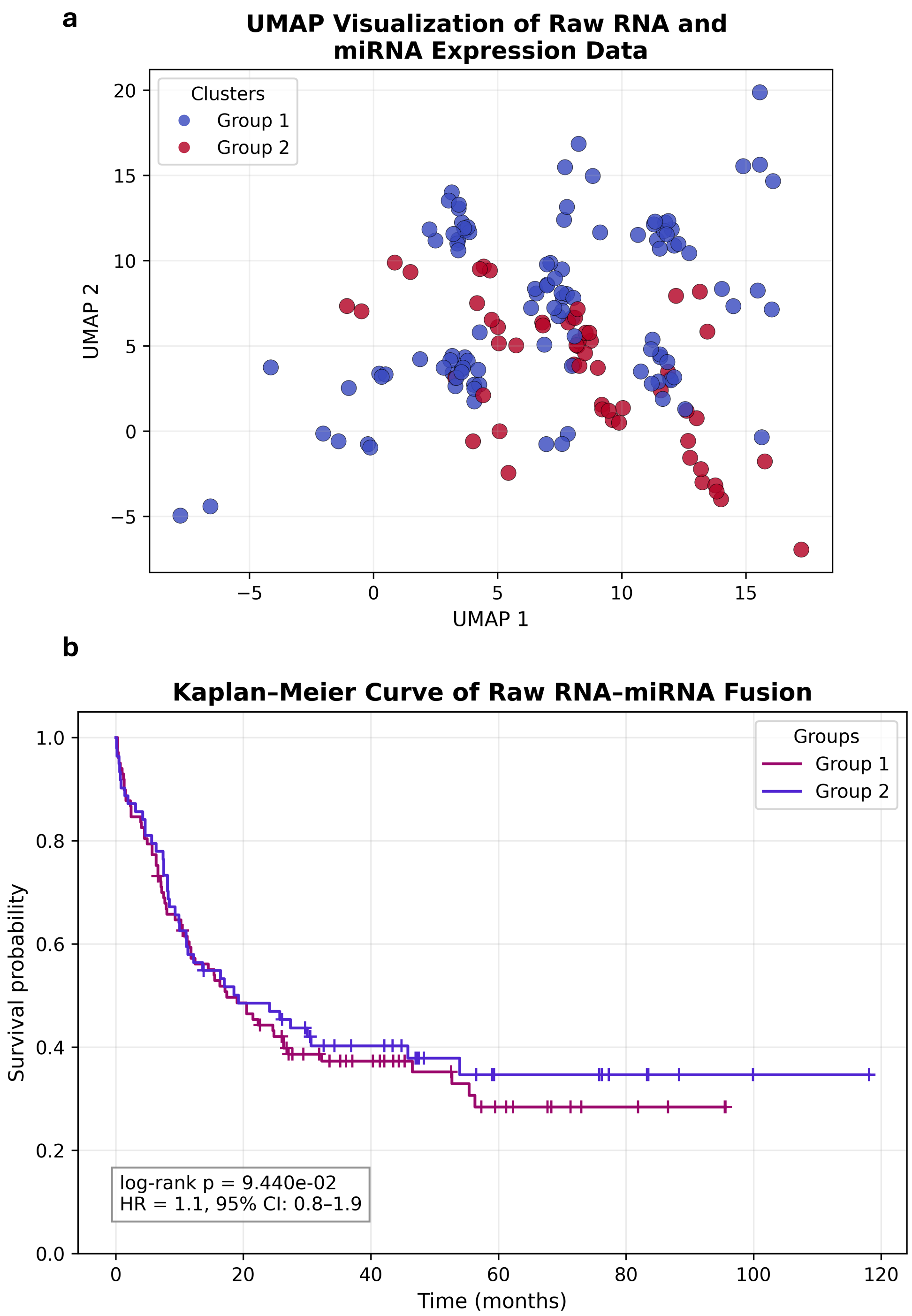

**Supplementary Fig 3.**

**Kaplan–Meier survival analysis of the concatenation-based clinical integration model.**

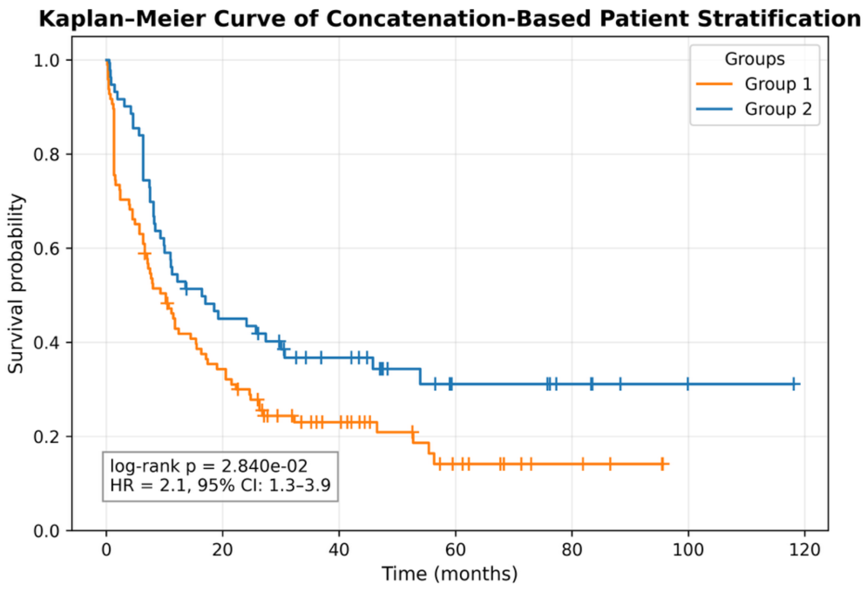

**Supplementary Fig 4.**

**Cross-modality reconstruction loss for single-omics models.**

(a) Reconstruction loss of gene-only and miRNA-only models for RNA and miRNA modalities. (b) Hazard ratios (95% confidence intervals) for model-derived patient groups obtained from the corresponding single-modality latent representations.

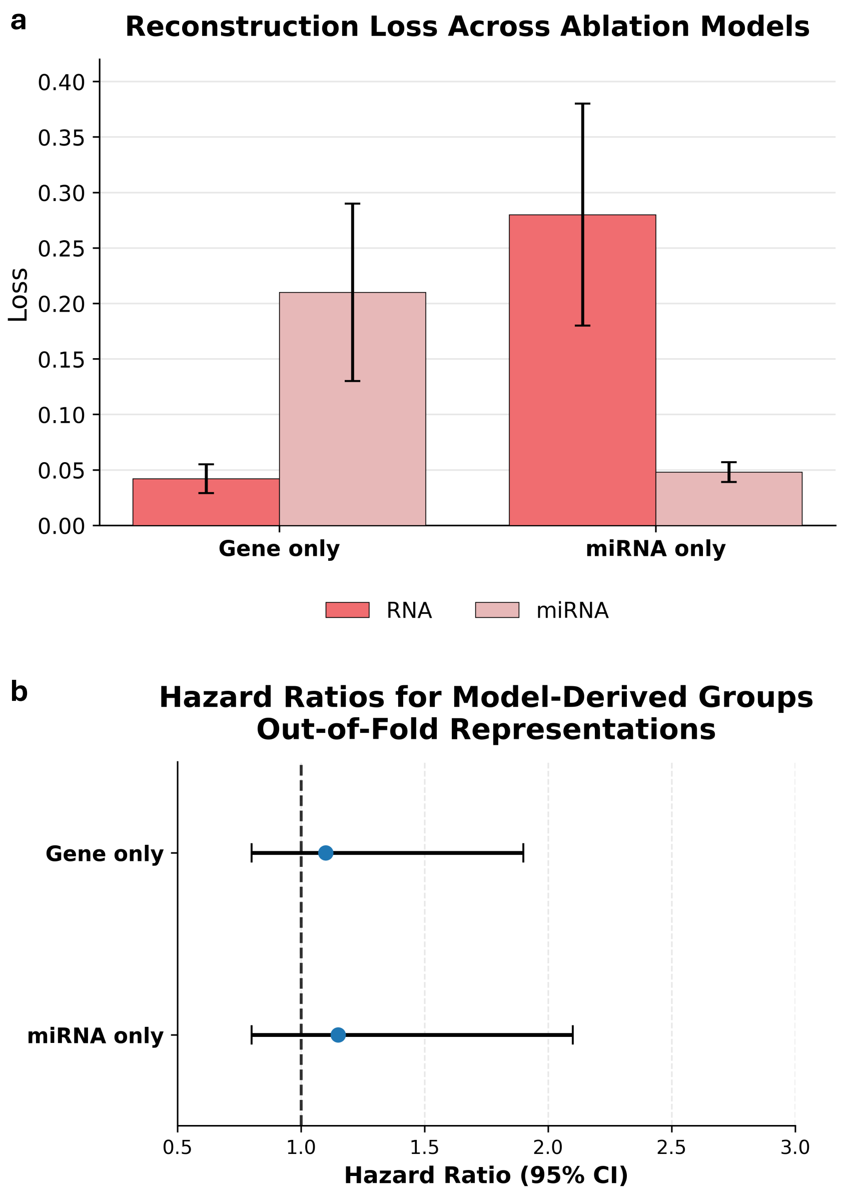

**Supplementary Fig 5.**

**Baseline multi-omics autoencoder model architecture.**

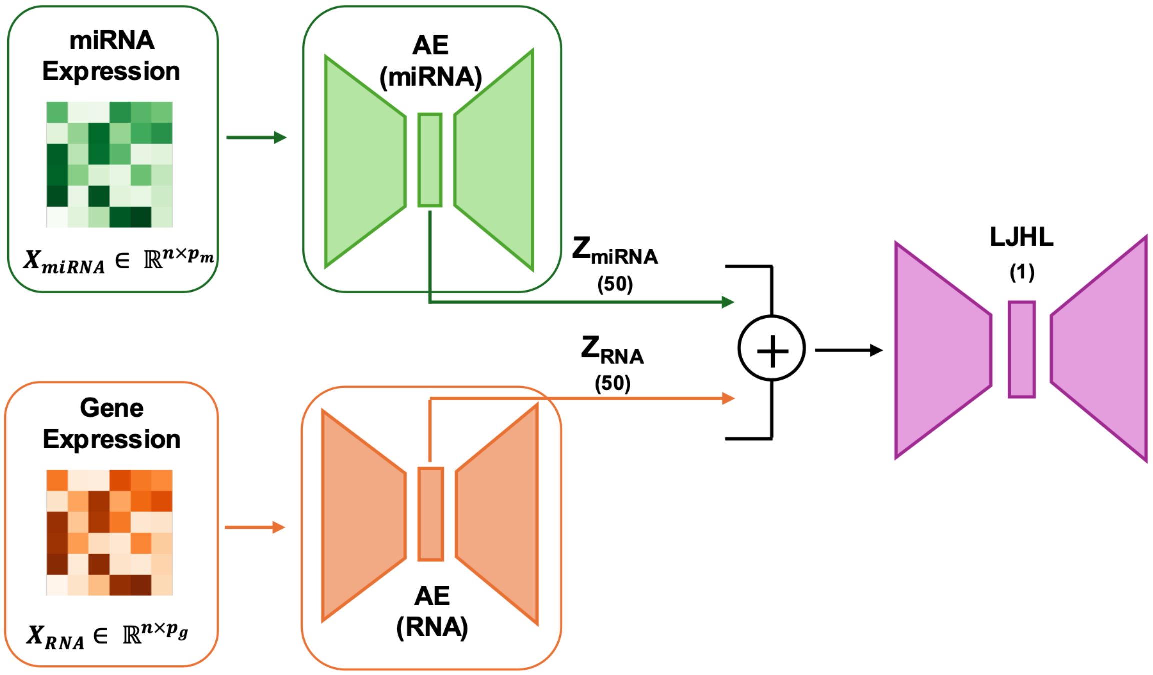

**Supplementary Fig 6.**

**Comparison of autoencoder and variational autoencoder models.**

Evaluation of baseline autoencoder (AE) and variational autoencoder (VAE) architectures. (a) Reconstruction correlation. (b) Reconstruction loss. (c) Survival stratification assessed by −log10(log-rank *p* value). (d) Hazard ratios (95% confidence intervals) for model-derived patient groups.

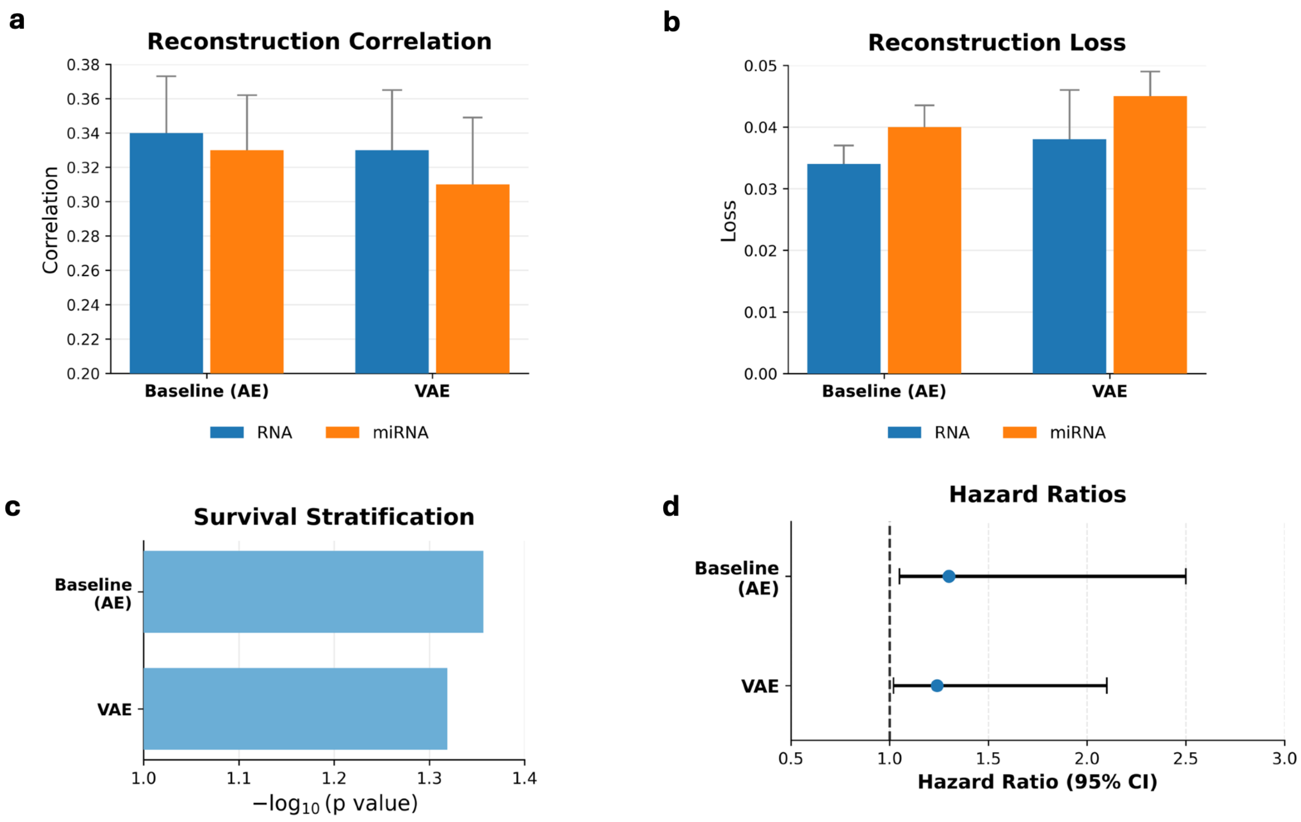

**Supplementary Fig 7.**

**Self-attention placement strategies evaluated.**

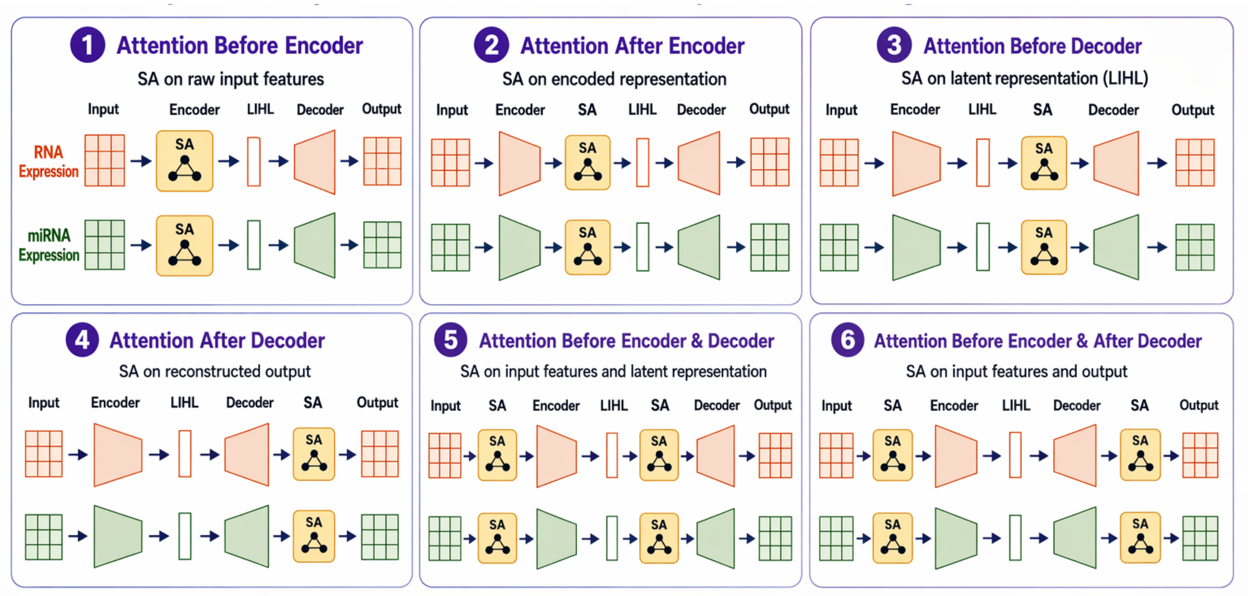

**Supplementary Fig 8.**

**Reconstruction loss across self-attention placements.**

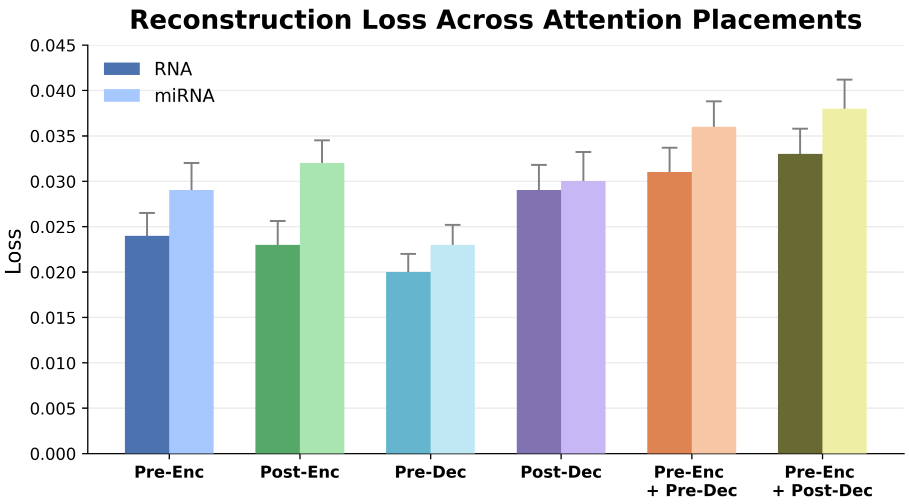

**Supplementary Fig. 9.**

**Reconstruction loss following the incremental incorporation of architectural components.**

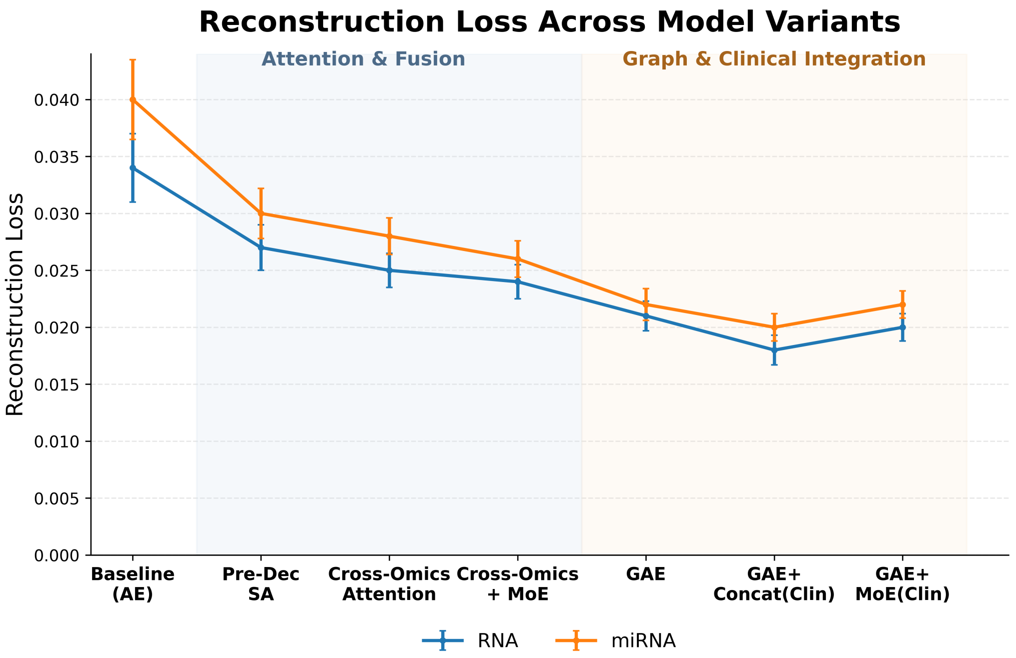

**Supplementary Fig. 10.**

**Hazard ratios following the incremental incorporation of architectural components.**

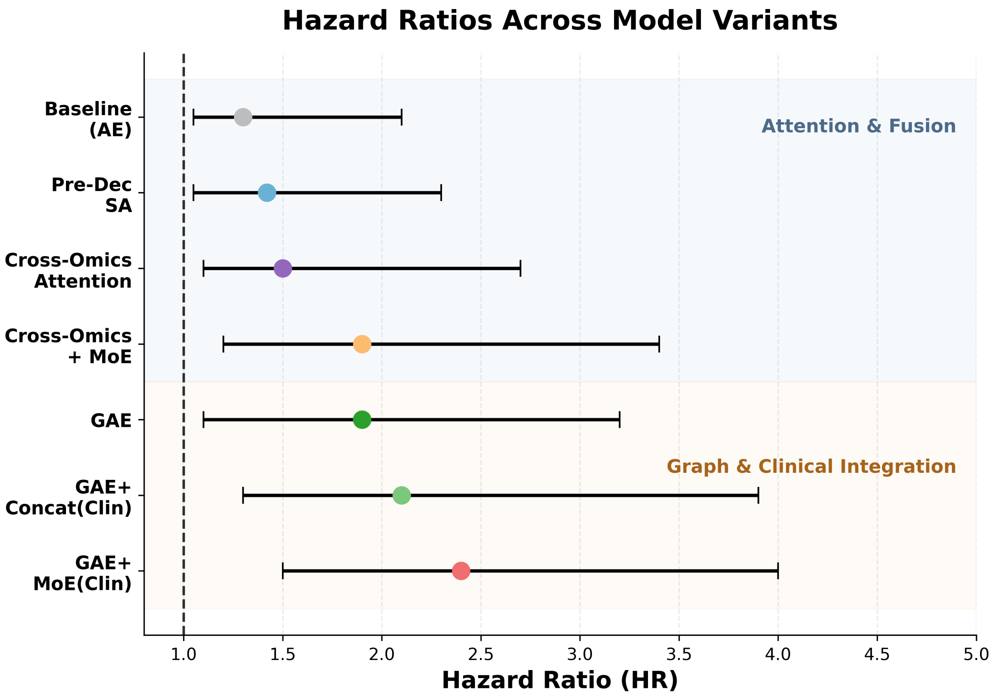

**Supplementary Fig 11.**

**Comparison of latent-space organization across multi-omics integration methods.**

UMAP projections are shown for (a) MO-GCAN, (b) CrossAttOmics, (c) MOFA-NC, and (d) MOFA-C.

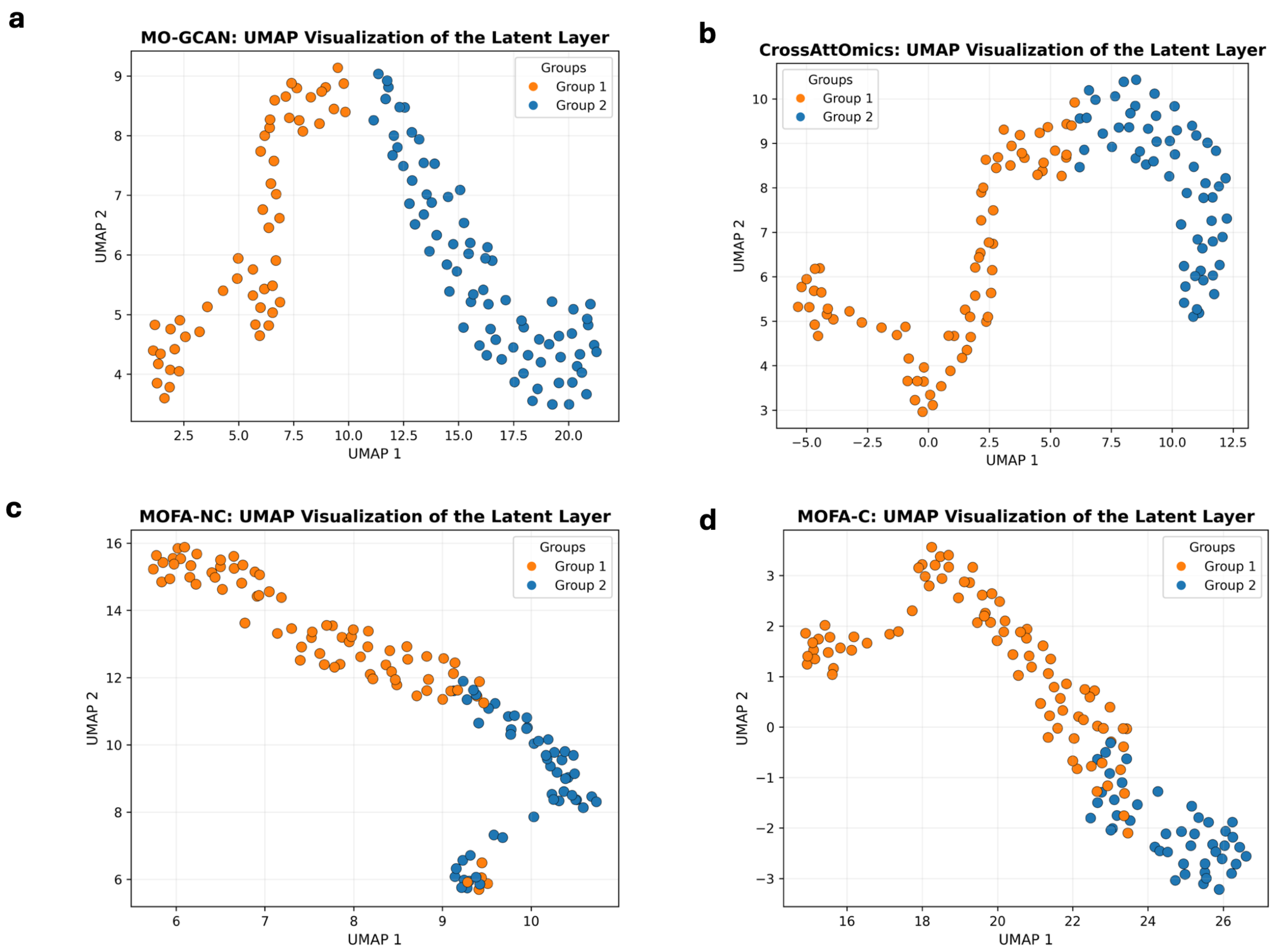

**Supplementary Fig 12.**

**Comparison of survival stratification performance across multi-omics integration methods.**

(a) Hazard ratios (95% confidence intervals) for model-derived patient groups. (b–e) Kaplan–Meier survival curves generated from out-of-fold latent representations for MO-GCAN, CrossAttOmics, MOFA-NC, and MOFA-C.

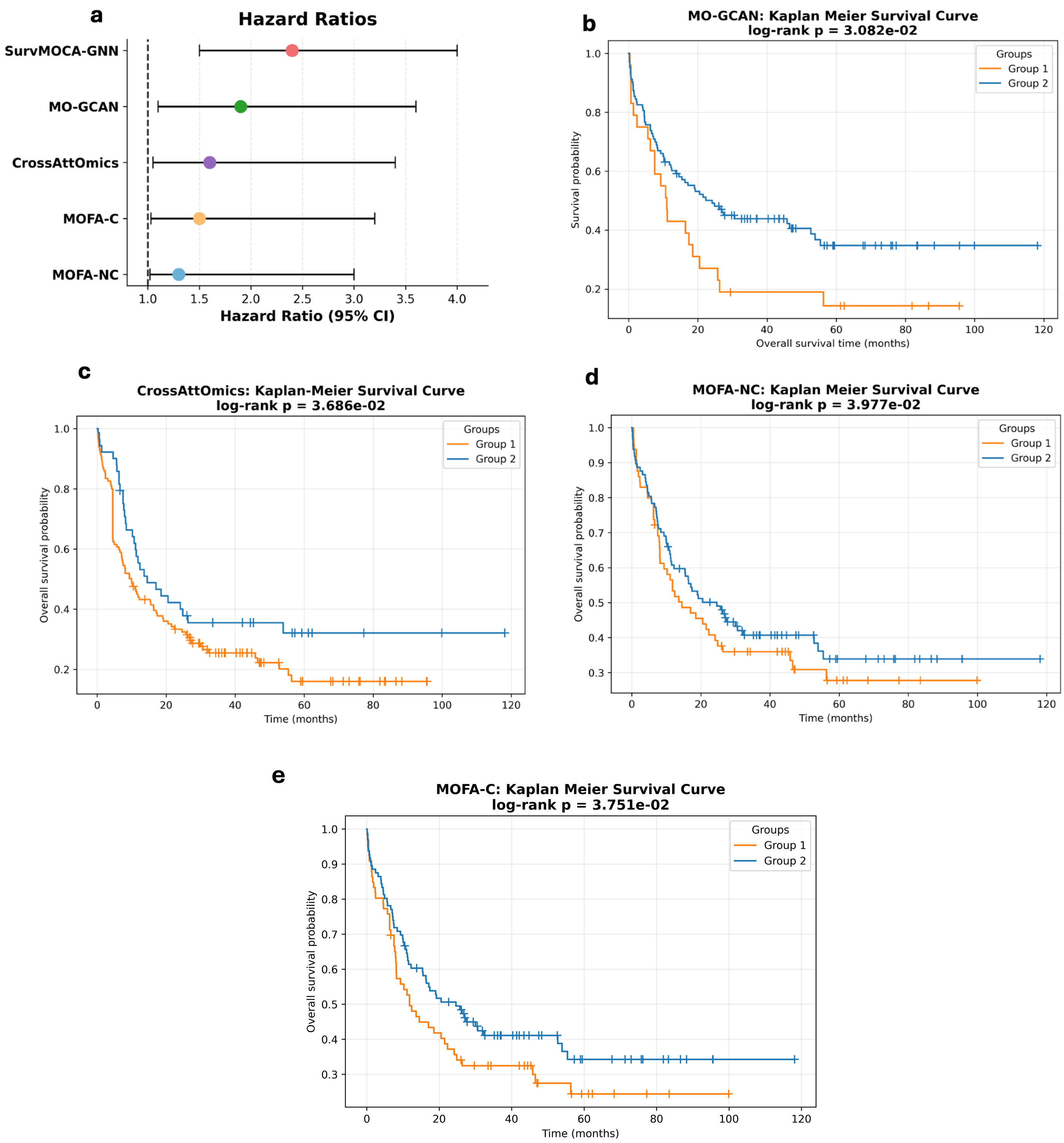

**Supplementary Fig 13.**

**External validation reconstruction loss of RNA and miRNA modalities on the external GAML cohort using direct validation of TCGA-trained models and transfer learning.**

Reconstruction loss for RNA and miRNA following direct application of TCGA-trained models and transfer learning using (a) MoE fusion and (b) concatenation-based integration.

**a**

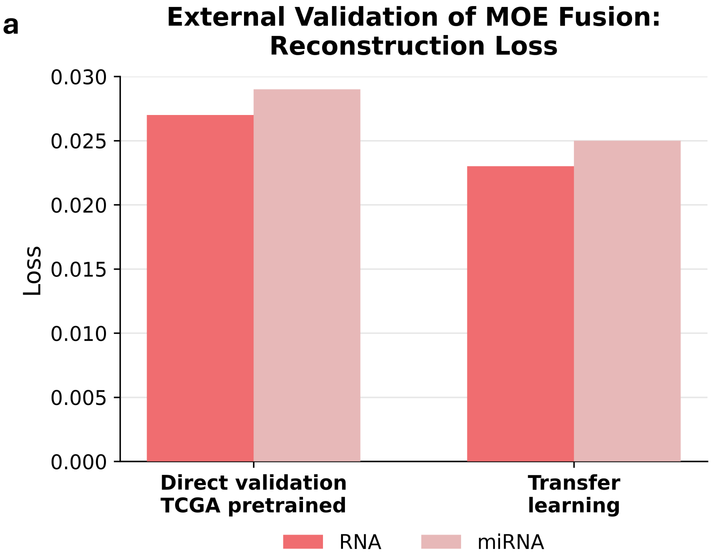


**b**

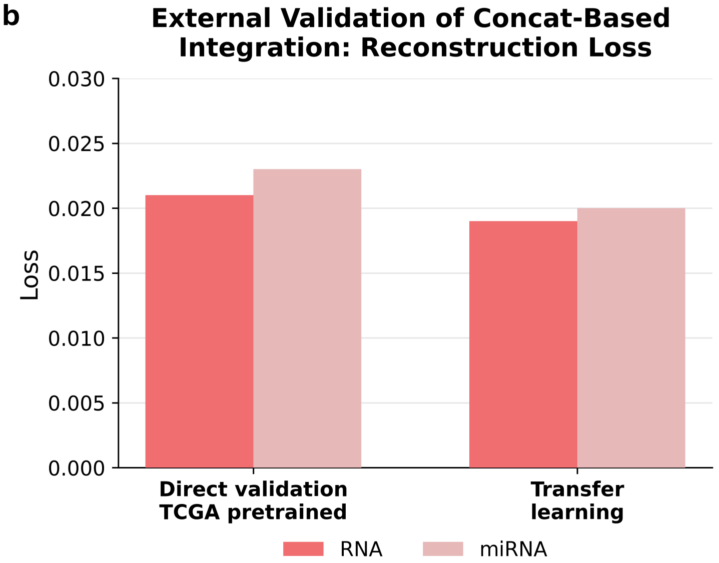

**Supplementary Fig 14.**

**Generalizability of learned latent representations in the independent GAML cohort.**

UMAP visualization of the learned latent representations in the external GAML cohort following (a–b) direct application of the pretrained model and (c–d) transfer learning. Panels show model-derived groups (a, c) and tumor/normal annotations (b, d).

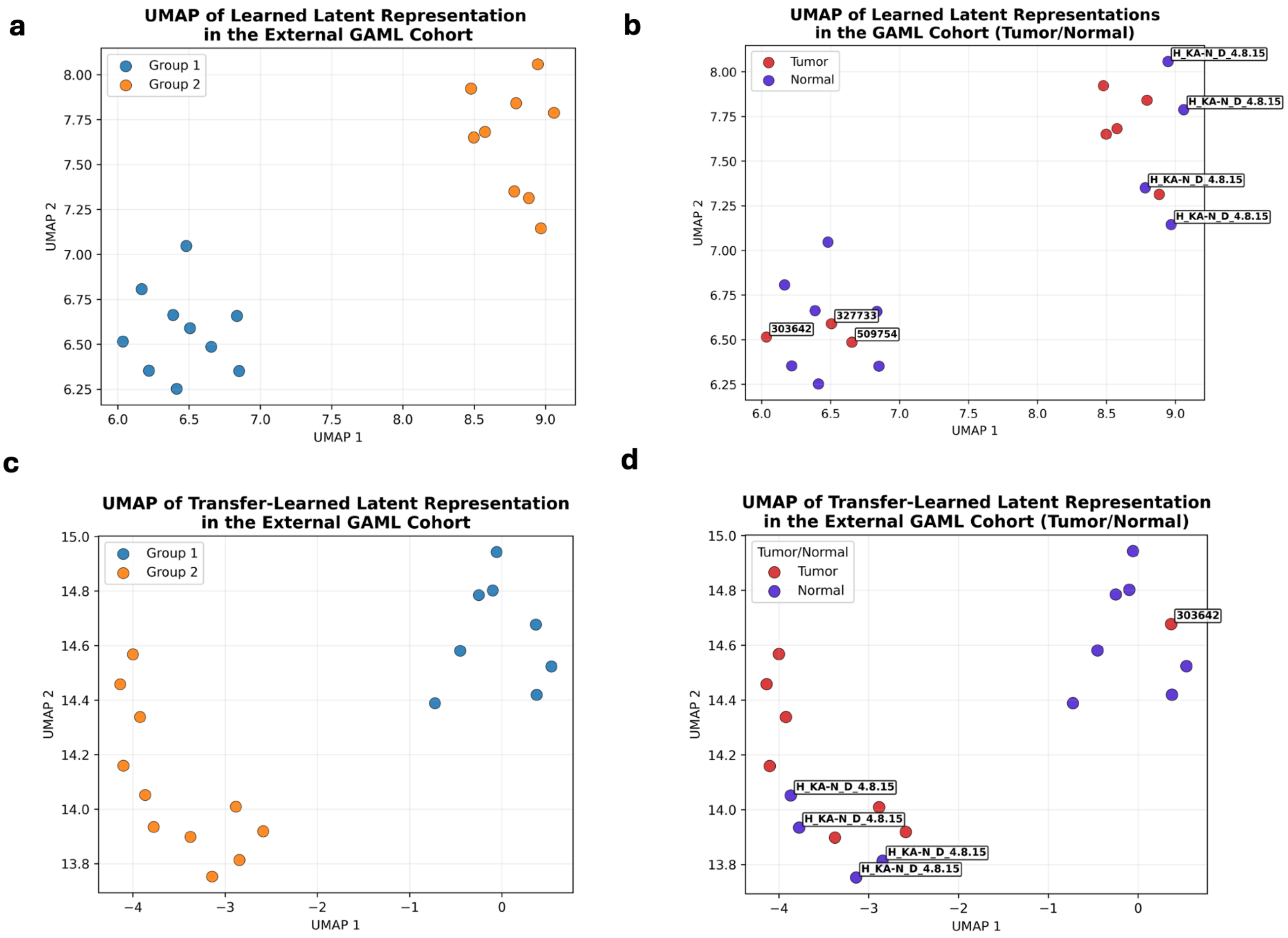
